# Systematic determination of in vitro phenotypic resistance to HIV-1 integrase strand transfer inhibitors from clinical samples

**DOI:** 10.1101/621755

**Authors:** Aniqa Shahid, Wendy W. Zhang, Vincent Montoya, Peter K. Cheung, Natalia Oliveira, Manraj S. Sidhu, Conan K. Woods, Marjorie A. Robbins, Chanson J. Brumme, P. Richard Harrigan

## Abstract

Phenotypic resistance data is relatively sparse for the newest HIV-1 integrase strand transfer inhibitors (INSTIs), dolutegravir (DTG), bictegravir (BIC), and cabotegravir (CAB). In this study, we report the phenotypic susceptibility of a large panel of oligo-clonal patient-derived HIV-1 integrase viruses. Representative clinical samples (N=141) were selected from a large database (N=17,197) of clinically-derived HIV integrase sequences, based on the presence of permutations of substitutions at 27 pre-defined positions in integrase (N=288). HIV-1 RNA was extracted from patient samples and diluted to approximately 500 HIV RNA copies/mL. Using an “oligo-clonal” amplification approach to achieve single-copy amplification, these dilutions were subjected to 12 parallel RT-PCR reactions to amplify integrase. Confirmed clonal amplicons were co-transfected with linearized pNL4.3∆int into CEM-GXR cells. In total, 162 HIV-1 viruses that carried no mixtures and had a unique sequence were harvested, and phenotyped in MT4-LTR-EGFP cells subsequently. Variants with the highest fold change (FC) had G140S and Q148R/H and resistant to all five drugs; R263K was the only single variant conferring >3-FC to DTG, BIC and CAB. There was extensive cross-resistance between DTG, BIC, and CAB and phenotypic resistance values for all the three INSTIs were almost collinear. The greatest exceptions were variants with N155H/G163E or L74I/T97M/F121C/V151I/E157Q/G163K, where both had >70-FC for CAB, while <3-FC for DTG and BIC. While site-directed mutagenesis is invaluable; the systematic selection of representative mutational patterns observed *in vivo* provides an efficient way to identify clinically relevant drug resistance.

## BACKGROUND

The success of combination antiretroviral therapy (cART) has remarkably reduced HIV/AIDS-related morbidity and mortality [1]. However, the emergence of antiretroviral (ARV) drug resistance can compromise global HIV/AIDS treatment and prevention strategies [2, 3]. With the increasing use of ARV drugs, HIV drug resistance (HIVDR) surveillance is warranted. Among other ARV drugs, integrase strand transfer inhibitors (INTSIs) are increasingly prescribed because of their tolerability and efficacy [4–6]. In particular, raltegravir (RAL), elvitegravir (EVG), dolutegravir (DTG), and bictegravir (BIC), are currently in clinical use [7], with cabotegravir (CAB) in advanced clinical development [8]. However, similar to other ARV drugs, emerging INSTI resistance may become a barrier to HIV disease management [9].

HIV drug resistance can be inferred via automated sequence-based genotyping, with interpretations of nucleic acids driven at least in part, by genotype-phenotype correlations. Genotyping is usually preferred over quantitative phenotyping for routine clinical testing because of lower costs, faster turn-around time, and greater accessibility [10]. However, phenotyping can provide insight required for adequate understanding of ARV drug resistance. Most importantly, it has been challenging to understand INSTI resistance because: a) phenotypic resistance data is relatively sparse for INSTIs; b) there are low reported failure rates of patients on INSTIs; c) *in vitro* drug resistance selection has been slow or unsuccessful [11]; and d) multiple mutations may be required to confer resistance [12].

While *in vitro* INSTI drug selection experiments, typically starting from clonal HIV subtype B isolates, have been instrumental in providing detailed insights to the emergence of novel HIVDR mutations [13]; these cannot represent the majority of virus sequence diversity observed *in vivo*. In order to capture the breadth of the naturally occurring viral variation in INSTI-naïve and experienced populations, it is essential to generate patient-derived integrase isolates and perform *in vitro* phenotyping. Therefore, in this study, we performed a relatively large-scale phenotypic analysis of HIV drug resistance in clinically-derived isolates, focusing on all five INSTIs to maximize coverage of *in vivo* sequence variation.

## MATERIALS AND METHODS

### Research ethics approval

The University of British Columbia Providence Health Care Research Ethics Board approved this research study (protocol H14-03423).

### Selection of clinical samples

The materials and methods employed in this study were previously reported [12]. The British Columbia Centre for Excellence in HIV/AIDS (BCCfE) routinely conducts clinical HIV genotypic drug resistance testing on total nucleic acid extracts (RNA/DNA) derived from patient plasma samples, and the DNA sequence data is regularly uploaded to the BCCfE’s secure Oracle database. For the purposes of this study, clinical samples with integrase genotypic resistance tests and their DNA sequences were identified from the BCCfE database (N= 17,197). A total of 27 amino acid positions in integrase were identified as having amino acids that differed in prevalence between INSTI-treated (primarily RAL and/or EVG) and naïve individuals, (https://hivdb.stanford.edu/cgi-bin/MutPrevBySubtypeRx.cgi) at the time of study design. Of the 27 amino acid positions, (51, 66, 68, 74, 92, 95, 97, 114, 118, 121, 128,138, 140, 143, 145, 146, 147, 148, 151, 153, 155, 157, 163, 230 and 263), 25 have been previously reported and are part of the Standard HIVdb ver. 8.4 interpretive algorithm for integrase (Supplemental Figure 1) [14, 15]. By comparing the odds (the frequency of mutation in INSTI-experienced individuals divided by the frequency of mutation in INSTI-naïve individuals) of mutations occurring at the positions along the integrase, positions 79 and 232 were identified as positions having high odds of mutations occurring in INSTI-experienced patients.

Having identified 27 codon positions, we next sought to identify clinical isolates harboring permutations of one or more of these substitutions *in vivo*. Clinical samples with mixtures at any of the 27 amino acids were excluded from further analysis and remaining 12,109 samples were categorized based on their 27 amino acid permutations. In total, 288 unique permutations of substitutions at the 27 codons were identified. For each permutation, one to three samples were selected for further processing, leading to a total of N=141 tested samples focusing on the most commonly occurring permutations which existed *in vivo*.

### Cell lines and reagents

CEM-GXR cells and HIV-1 NL4.3 plasmid with integrase gene deleted (pNL4.3∆int) were generously provided by Dr. Mark Brockman (Simon Fraser University, Burnaby, Canada) [16]. Dr. Theresa Pattery (Janssen Diagnostics, Beerse, Belgium) kindly provided MT4-LTR-EGFP cells. The integrase inhibitors, RAL, EVG, DTG, and CAB were purchased from Selleck Chemicals (Houston, USA). Bictegravir was synthesized by the Centre for Organic Synthesis, University of British Columbia (Vancouver, Canada).

### Oligo-clonal amplification and sequencing

Total nucleic acids were extracted from 500µL of plasma using the NucliSens easyMAG Extractor (bioMérieux, Saint-Laurent, Canada). For most samples tested, HIV RNA from routine clinical genotyping was stored at −80°C. If RNA was not available after routine clinical genotyping, a fresh extraction was performed. Oligo-clonal amplification was performed on total nucleic acid extracts by diluting them in diethyl pyrocarbonate (DEPC)-treated water to approximately 250-500 HIV RNA copies/mL to achieve single-copy amplification. Diluted extracts were amplified in 12 parallel reactions using the Transcriptor One-step RT-PCR kit (Roche, Basel, Switzerland) for reverse-transcription and first-round PCR. A second round of PCR amplification was performed on each reaction using integrase-specific primers, generating an amplicon that covered 1-288 amino acids of integrase [12]. PCR amplicons were sequenced on an ABI 3730×l sequencer (Thermo Fisher Scientific, Waltham, USA) and analyzed using the in-house base-calling software ReCall (University of British Columbia, Vancouver, Canada) [17]. These conditions led to about a 50% success rate for PCR amplifications; assuming a Poisson distribution, most amplifications would have originated from 0, 1 or 2 original starting templates. Amplicons that contained no inferred amino acid mixtures at 27 codons associated with HIV INSTI exposure upon sequence analysis were chosen for recombinant virus production in order to have relatively direct genotype-phenotype correlations.

### Recombinant virus production

Linearized pNL4.3∆int was generated using BstEII restriction digestion as previously described [16]. Recombinant viruses were generated by co-transfecting linearized pNL4.3∆int with the corresponding second round PCR product into CEM-GXR cells, via electroporation using a Bio-Rad GenePulser II. Transfected cells were transferred into tissue culture flasks with 5 mL of room temperature Roswell Park Memorial Institute (RPMI) culture medium supplemented with 20% fetal bovine serum (R20+) and incubated at 37 °C in a humidified 5% CO_2_ atmosphere. After co-transfection, 2 mL of fresh R20+ was added every two to three days until the total volume of each tissue culture flask reached 10 mL. After which, 2 mL of supernatant was removed and replenished with fresh R20+ media until harvest. Green fluorescent protein (GFP) expression was monitored by flow cytometry (FACSCalibur Guava 8HT [Millipore]) starting on day 12, as previously described [16]. Once the infection (GFP-positive cells) reached 15-30%, viruses were harvested and stored at −80°C until use. HIV-1 RNA was extracted from the harvested viruses and subjected to nested RT-PCR and Sanger sequencing of integrase, as described previously [12, 17]. The harvested recombinant virus sequences were confirmed to be identical to the amplicon sequences at 27 codon positions. Recombinant viruses with a unique permutation of amino acids at the 27 positions of interest were assayed for phenotypic drug resistance. For viruses, which were identical at the 27 positions of interest, only a single virus (chosen arbitrarily) was assayed.

### Phenotypic drug susceptibility assays

Recombinant virus titres were determined by infecting MT4-LTR-EGFP cells in a three-day assay and infectivity data collected using a SpectraMax i3 Minimax 300 microplate reader and cytometer (Molecular Devices, San Jose, USA). The SpectraMax captures images of each individual well of a 96-well plate and counts the number of infected (GFP-positive cells) and non-infected (GFP-negative) cells. Titered volumes of recombinant viruses (such that 15% - 30% infection is reached on the third or fourth day post-infection) and MT4-LTR-EGFP cells were plated in triplicate with eight concentrations ranging from no drug to 1000nM (10-fold dilutions from 0.1nM to 1000nM with 3nM and 45nM included) of BIC, CAB, DTG, RAL, and EVG. Viral infectivity data was measured using the SpectraMax microplate reader

### Data analysis and graphing

Graphs and statistical analysis were done in R (version 3.2.3). The 50% effective concentrations (EC50s) were calculated by fitting to a four parameter EC50 model using in-house scripts written in R using the nplr package version 0.1-7. Fold changes (FC) in EC50 of the virus relative to a NL4.3 control virus were calculated using mean EC50s (performed in triplicate). Fold change values were log transformed and presented as log-FC.

## RESULTS

### Systematic determination of amino acid permutations to test in vitro

To maximize the efficiency of HIV integrase phenotyping, we implemented a systematic approach to identify clinical samples capturing the most commonly occurring amino acid permutations at 27 positions in integrase associated with INSTI exposure *in vivo*. A total of 288 amino acid permutations at these positions were observed in our large clinical database (Supplemental Table 1). To maximize the direct relationship between genotype and phenotype correlations, these samples were further diluted and amplified in 12 parallel RT-PCR reactions to generate near clonal DNA sequences (see Methods) to minimize confounding by “mixtures”. The overall PCR amplification rate that led to a successful amplification of the integrase gene with no mixtures at the 27 inferred amino acid positions averaged approximately 41% (1644/3985 amplicons, including repeated tests). A total of 162 recombinant viruses were generated and phenotyped. Of note here, all recombinant viruses (N=162) were phenotyped for DTG, BIC and CAB. However, a subset of these (N=84) were phenotyped for RAL and EVG (Supplemental Table 3). These viruses represent 90% of the observed sequence variation among the clinical samples at the 27 relevant integrase codon positions. The results from the 141 clinical samples representing six HIV-1 subtypes (B=101, C=16, A=9, G=3, D=2, F=1) and 3 circulating recombinant forms (CRF02_AG=4, CRF01_AE=4, CRF03_AB=1) from the BCCfE database are presented in this study (Supplemental Tables 2 and 3), indicating both the complete amino acid sequence and the most relevant 27 codon positions. At the 27 codons, the viruses (N=162) carried up to 50 amino acid substitutions relative to reference HIV subtype B consensus sequence (Figure 1).

**Figure 1.**
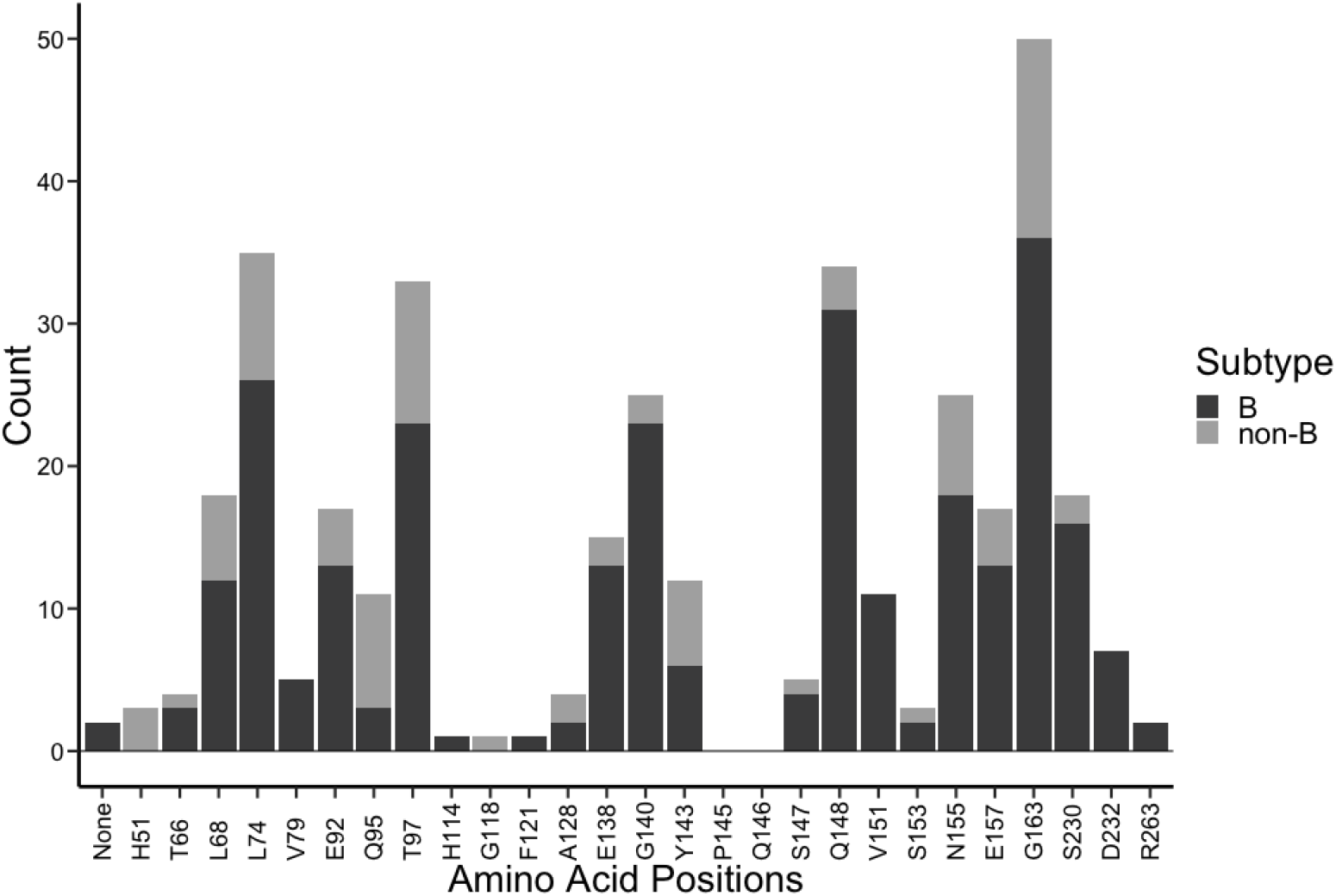
The number of samples with amino acid substitutions observed at each of the 27 integrase codons investigated in subtype B (N=115, dark grey) and non-B viruses (N=47, light grey).

### Differential selection of resistance to INSTIs

We assayed 162 recombinant viruses for phenotypic susceptibility to five INSTIs; raltegravir, elvitegravir, dolutegravir, bictegravir and cabotegravir. All amino acid substitution combinations and their observed FC values for the five INSTIs are listed in Supplemental Table 3. We observed a wider log-FC shift for subtype B compared to non-B viruses (Figure 2). Among single variants, R263K was the only single variant conferring >3-FC for all DTG, BIC and CAB (3.9, 3.2 and 4-fold, respectively). The single mutants N155H, E157Q and Y143R had >3-FC for RAL and EVG. Among the double mutants observed, T66I/G118R had >3-FC for DTG, BIC and CAB. G140S/Q148R showed consistently >3-FC for all five drugs. N155H/G163E had >3-FC for CAB (and RAL and EVG) but was susceptible to DTG and BIC. R263K/G163E had consistently higher FC (>3-FC) as a double mutant as well. E138K/Q148R had >10 and >200-FC for RAL and EVG, respectively. Among triple mutants observed, E138A/G140S/Q148H showed >5-FC for CAB, however it was susceptible to BIC and DTG (2 and 2.8-FC, respectively). As shown previously [12], eight viruses carried G140S/Q148H only, two had the combination G140S/Q148H + T97A and 3 viruses had G140S/Q148H + T97A + L74M. Viruses with G140S/Q148H were highly resistant to RAL and EVG with >100-FC but had relatively small changes in susceptibility to DTG, BIC and CAB (median 2.9 to 4.7 FC). Viruses with an additional T97A substitution or additional T97A + L74M substitutions maintained high FC to RAL and EVG (>100-FC) but were increasingly resistant to DTG, BIC, and CAB [12]. Among quadruple mutants, V79I/E138K/G140A/Q148R had >10-FC for DTG, BIC and CAB. A virus carrying six amino acid substitutions (L74I/V79I/G140A/Q148R/V151I/ E157Q) showed high FC to DTG, BIC and CAB (>500-FC). One additional virus carrying six amino acid substitutions (L74I/T97M/F121C/V151I/E157Q/G163K) showed >50-FC for CAB, however it was susceptible to DTG and BIC. The greatest exceptions were variants with N155H/G163E or L74I/T97M/F121C/V151I/E157Q/G163K, where both had >75-FC for CAB, while <3-FC for DTG and BIC (Figure 3).

**Figure 2.**
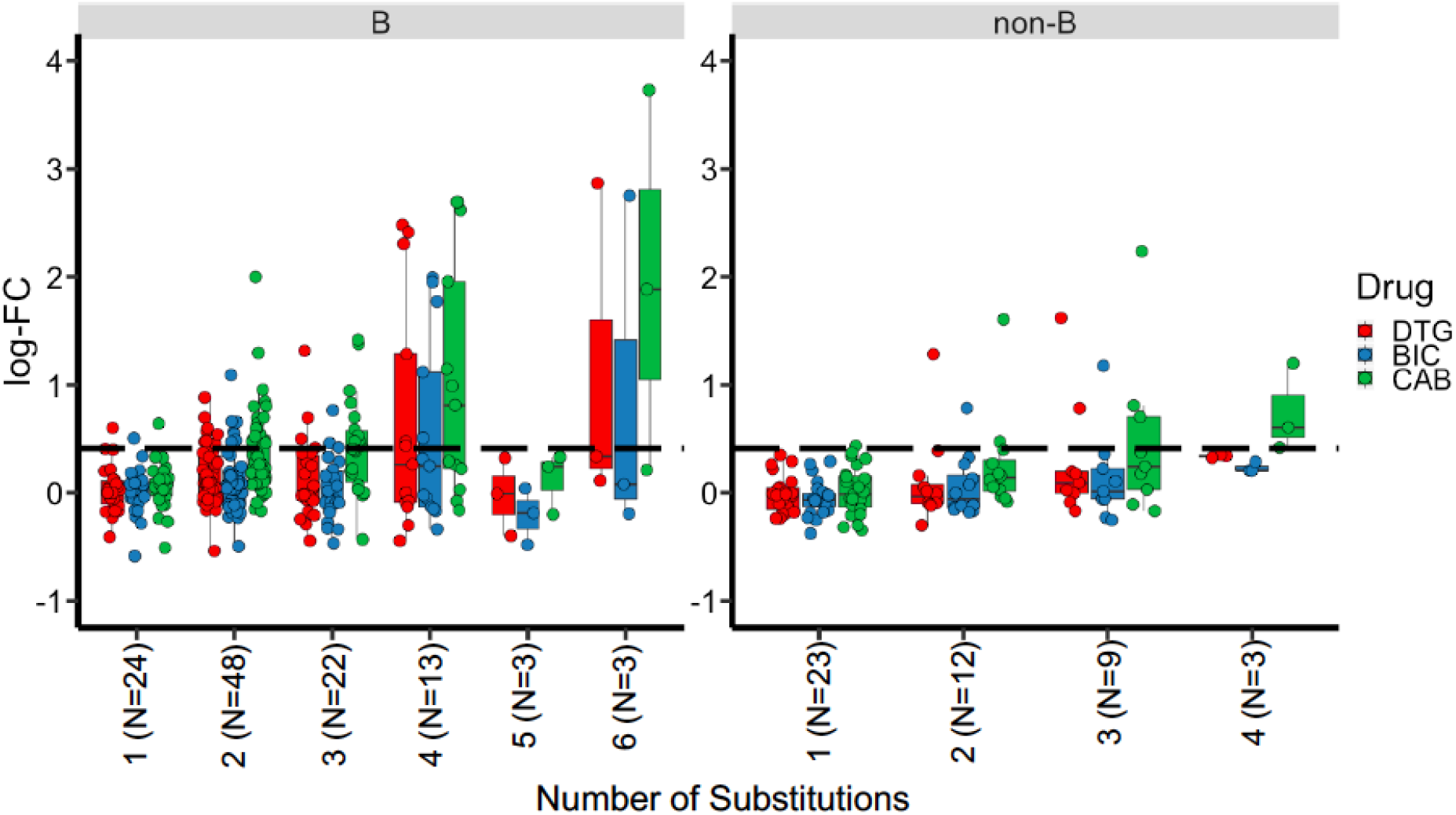
Log-fold change (FC) in DTG (red), BIC (blue), and CAB (green) of subtype B and non-B viruses with 1, 2, 3, 4, 5, and 6 amino acid substitutions relative to HIV subtype B consensus at 27 integrase codons of interest. The dashed line represents the PhenoSense integrase assay (Monogram Biosciences, Inc.) biological cutoff of log-transformed 2.5 FC for DTG as reference.

**Figure 3.**
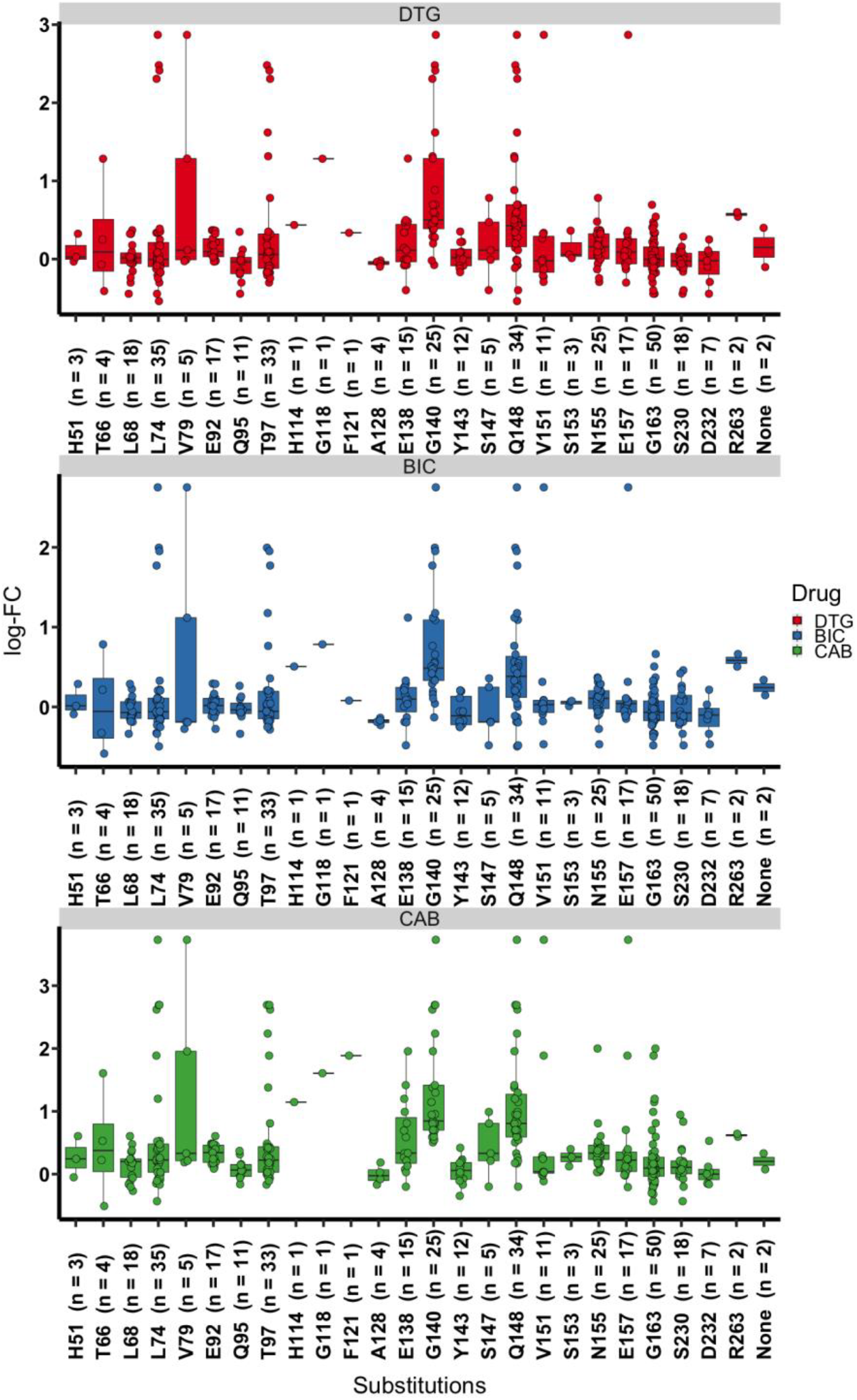
Log-fold change (FC) of each individual single amino acid substitutions in subtype B and non-B viruses.

### Extensive cross-resistance between DTG, BIC and CAB

Spearman rank correlation coefficients of log-FC values between DTG and BIC or CAB were strongly correlated, with BIC and CAB with correlation coefficients of 0.93 (slope 0.81) and 0.87 (slope 1.1), respectively (Figure 4A, 4B). DTG resistance, for select viruses where log-FC values were calculated for RAL and EVG, was modestly correlated with RAL and EVG resistance (data not shown), at least in part because RAL and EVG FC values exceeded what could be measured in our assay (>1000-fold).

**Figure 4.**
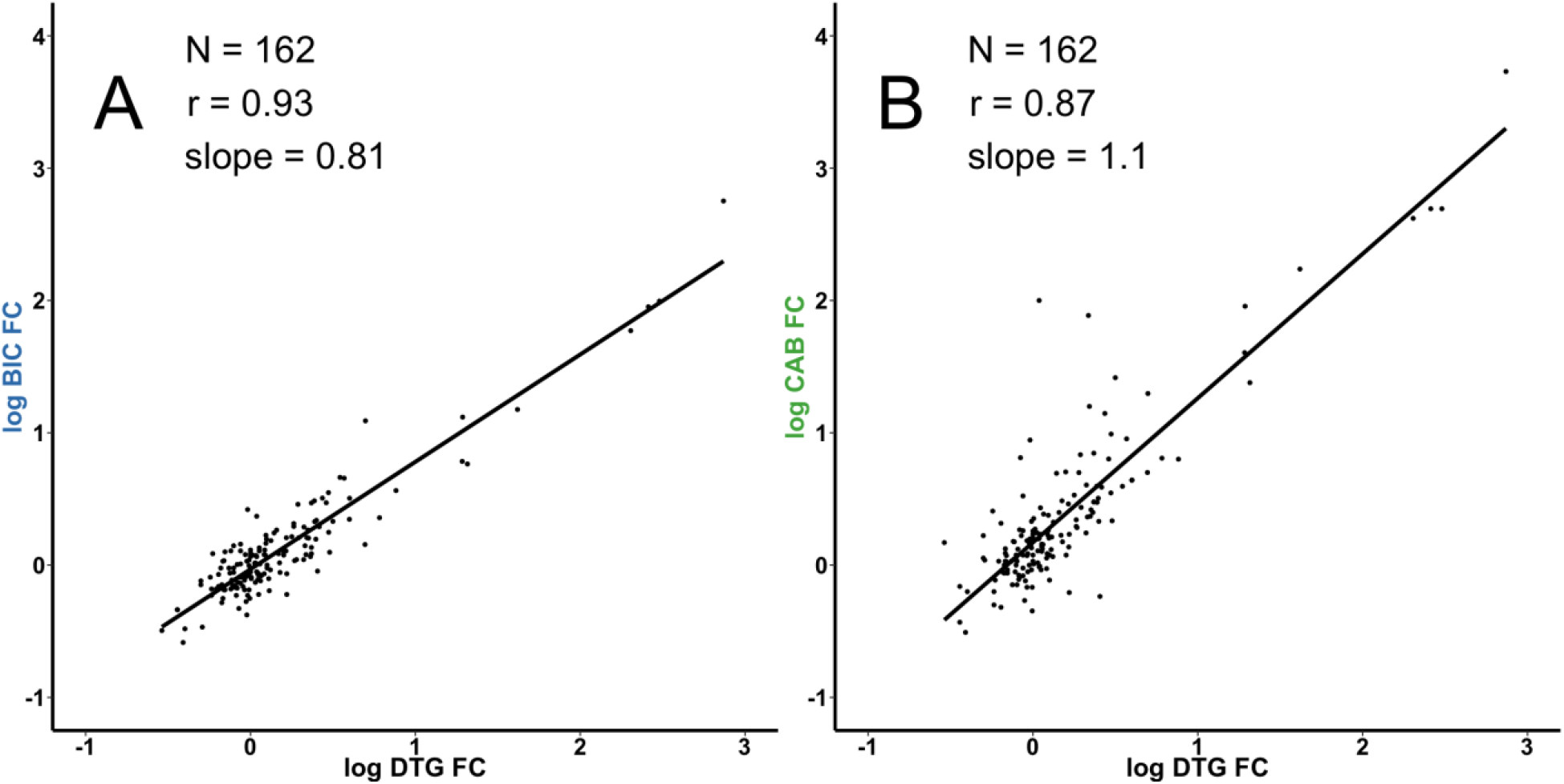
Correlation between log bictegravir (BIC), cabotegravir (CAB) and Dolutegravir (DTG) fold change (FC) values for N=162 viruses. A) Log BIC FC versus Log DTG FC. Correlation (r-value) was 0.93 and slope was 0.81. B) Log CAB FC versus Log DTG FC. Correlation (r-value) was 0.87 and slope was 1.1.

## DISCUSSION

The second-generation INSTIs, DTG and BIC have a higher genetic barrier to resistance in comparison to first-generation INSTIs (RAL and EVG), resulting in low rates of resistance development *in vivo* or *in vitro*. This desirable property has had the side-effect that the identification of drug resistance (especially by routine integrase sequencing methods) is difficult for second-generation INSTIs. However, this pressure may force HIV to evolve along novel and unidentified mutational pathways *in vivo*, including at sites outside the HIV *pol* gene [18]. Systematically screening large panels of clinically-derived isolates from patients offers insights into resistance phenotypes conferred by integrase variation and can inform genotypic resistance interpretation algorithms, such as the Stanford HIVdb [15].

The rationale behind our systematic screening strategy is worth the discussion here: 1) INSTI treatment failure reports are rare, especially for second-generation INSTIs [19]; 2) testing FC *in vitro* for all possible variations in integrase is an intractable problem; 3) in reality, only those mutations associated with INSTI Failure (or just INSTI exposure) require testing, as much of the other variation in integrase will not result in resistance - we therefore identified 27 codon positions that were associated with INSTI exposure by identifying mutations with greater prevalence in INSTI-treated patients in our database, as well as the data available in public domains (such as Stanford HIVdb); 4) in addition, integrase resistance may be the result of the accumulation of multiple resistance mutations *in vivo*, it is necessary to test combinations/permutations of mutations at these 27 codon positions; since this is impossible, even for 27 codon positions, we reduced this further by only testing permutations/combinations that were observed *in vivo*.

Theoretically, the numbers of mathematically possible permutations of the integrase, even considering only the 27 codons here vastly exceeds the number of permutations that were observed *in vivo* (N=288) from our database of >15,000 clinically-derived integrase sequences. In this way, relatively few samples can represent much of the observed *in vivo* variation (Supplemental Figure 2).

This is one of the first studies to report *in vitro* phenotypic susceptibility of a large panel of HIV-1 integrase recombinant viruses for DTG, BIC and CAB, derived directly from clinical samples where patients were primarily either integrase-naïve or exposed to first-generation INSTIs. Upon comparing our panel of viruses with published data (available on Stanford HIVdb (https://hivdb.stanford.edu/pages/phenoSummary/Pheno.INI.Simple.html), we observed that very few clinically-derived isolates resistant to DTG were reported previously. For DTG, 18 clinical variants have been reported to carry G140S/Q148H with 3 to 15-FC. Addition of T97A (G140S/Q148H/T97A) increased the resistance to >50-FC. This is broadly similar to what we have observed. However, there have been no reports on variants carrying L74M/T97A/G140S/Q148H. Moreover, there is no data on BIC and CAB other than our report here and some recent conference proceedings [20–22].

In addition, two variants containing G140S/Q148R showed modest FC values for DTG (4 and 6.2) [23, 24]. This is similar to our reported FC values of 3.7 and 4.9. There have been reports on laboratory-adapted strains that carry Q148R/N155H with DTG FC of about 5, but we have not observed a N155H mutation co-exist with a 148R in over 15,000 clinical isolates. Most commonly, N155H and Q148K exist in combination with G140S in our database. This suggests the difficulty of creating laboratory generated strains using site-directed mutagenesis that accurately represent the *in vivo* sequence observed in clinically-derived samples. In addition, we have not yet observed a variant carrying E138K/S147G/N155H/T97A/V151I, previously reported to be highly resistant to DTG (>50-FC) [25].

Consistent with previous reports, we observed that for the first-generation INSTIs RAL and EVG, higher log-FC values (exceeding limits of our analysis) for viruses carrying G140S and Q148H. However, these viruses were still susceptible to second-generation INSTIs DTG, BIC and an investigational drug, CAB. The addition of one or more substitutions (T97A and L74M) conferred greater resistance to DTG, BIC and CAB. Results were concordant with previously published data and confirmed extensive cross-resistance between DTG, BIC and CAB [12].

Some limitations of the study merit mention here. First, this study reflects populations either primarily exposed to RAL/EVG, or with no known prior exposure to INSTIs and as such, the >15,000 sequences considered here does not capture all sequence variation likely to be observed in patients failing second-line INSTI. Second, our focus on 27 amino acid substitutions only within the integrase gene may omit the contributions of other substitutions inside the integrase gene in conferring resistance, so we cannot make causal conclusions between the observed phenotypes and genotypes. Moreover, we have not looked at amino acid substitutions outside integrase which may confer low, medium and/or high-level resistance to the current INSTIs [18]. Third, we have not demonstrated any relevant cut-offs for inferring clinically relevant resistance and impact on virological outcomes. We recognize that the accurate interpretation of genotypic drug resistance data requires correlations with phenotypic data (*i.e*. genotype:phenotype relationships) as well as with clinical data on virological outcomes after therapy [26, 27].

In conclusion, naturally occurring amino acid variants emerging during ART provide insights into viral escape and resistance. Selection of patient-derived clonal viruses based on large genotype databases to make the observed sequence variation *in vivo* can be used to efficiently generate panels of resistant viruses for phenotype analysis. If new mutations or permutations are identified it is straightforward to select these for future phenotyping and identify new patterns leading to decreased susceptibility to the newest INSTIs. It will be essential to monitor HIV variation inside and outside the integrase gene in those patients who have failed DTG and/or BIC, particularly in those with non-B subtypes of HIV.

## Notes

### Acknowledgement

We acknowledge Sarina Barnes and Rob Hollebakken for technical assistance with PCR and sequencing. We thank Kimia Kamelian for critically reading the manuscript and providing a valuable feedback.

---

**Supplemental Figure 1.**
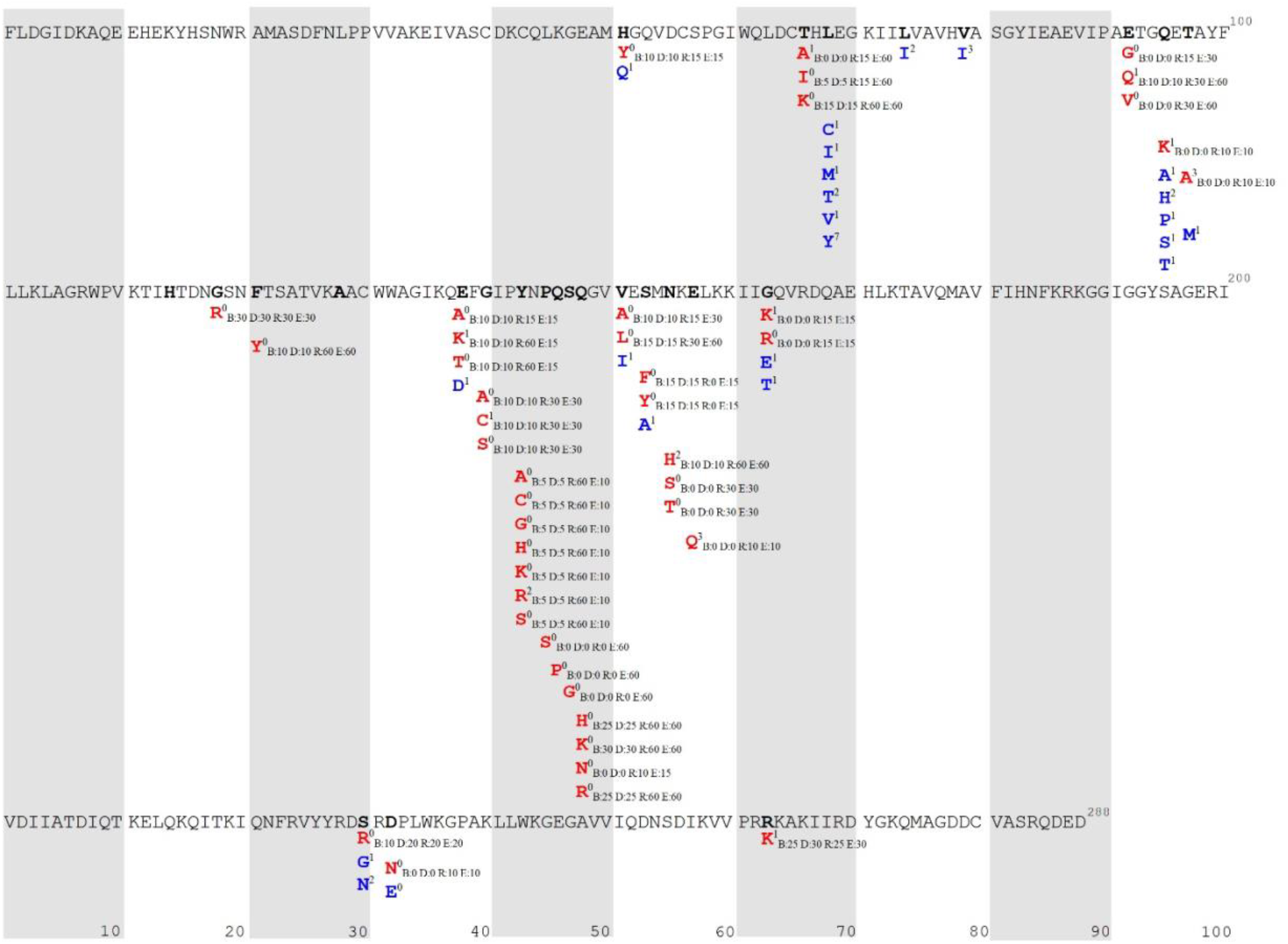
HIV-1 integrase map using subtype B consensus sequence as reference. Bolded black amino acids show 27 codon positions included in this study. Integrase inhibitor (INSTI) resistance positions and mutations are shown in red. Superscript number shows the frequency of amino acids observed in this study. Adjacent plain text indicates current Stanford INSTI resistance scores; no resistance [0] to high resistance [60] for bictegravir [B], dolutegravir [D], raltegravir [R], and elvitegravir [E]. Blue amino acids represent other variants observed at that particular codon, followed by its frequency.

**Supplemental Figure 2.**
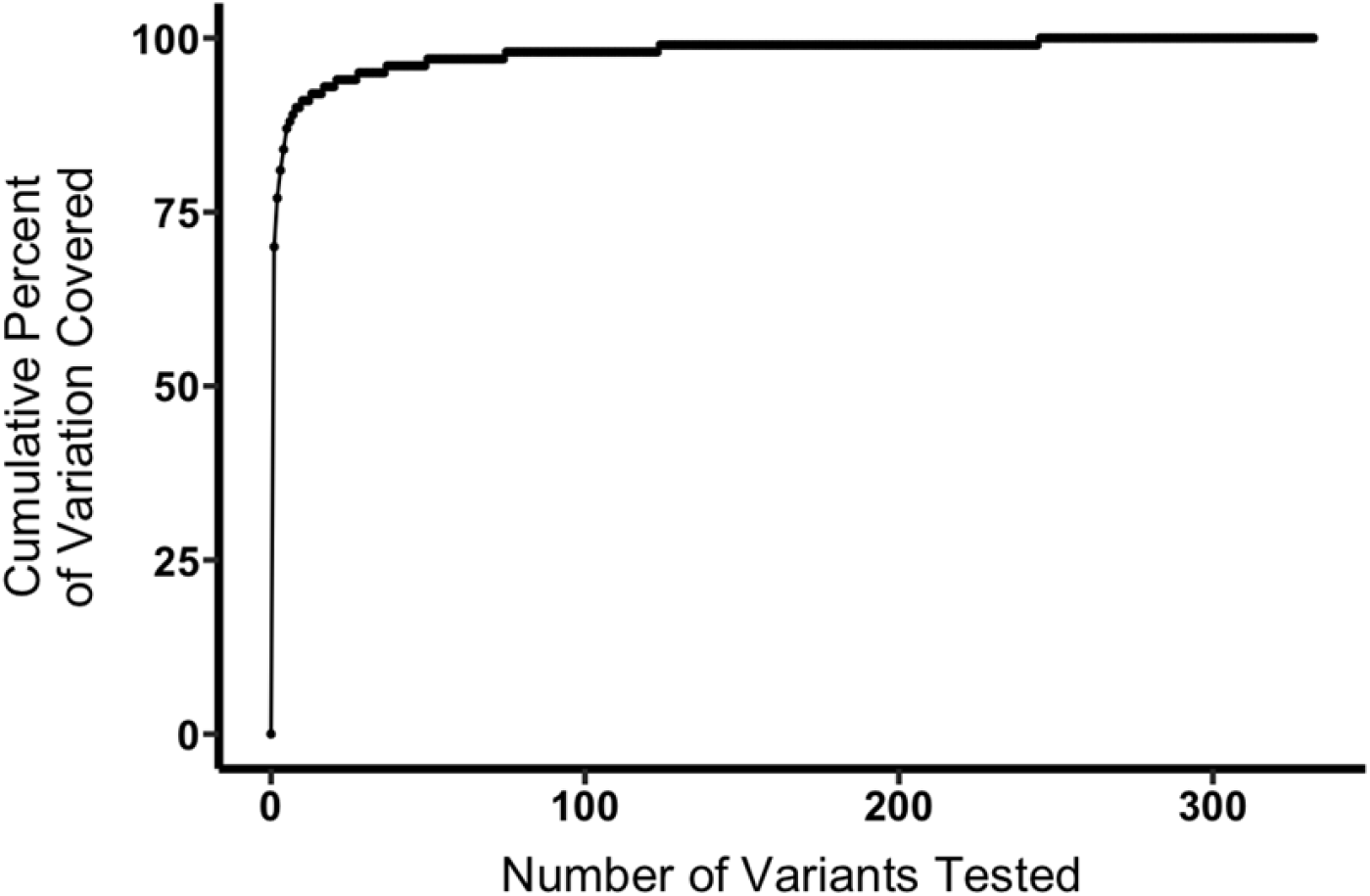
Number of variants tested from BCCfE database plotted versus their cumulative percentage shows that relatively few samples are required to represent much of the *in vivo* variation. Note that more sequence data were added to the existing 12,109 samples (as presented in Supplemental Table 1) later in the study, which increased the total number to 15,285 samples. The latter number was used to perform coverage analysis presented here.

**Supplemental Table 1.** This table lists the number of clinical samples to show known amino acid permutations (N=288) at 27 codon positions in the BCCfE clinical database.

**Supplemental Table 2.** This table lists all the patient-derived HIV-1 integrase recombinant viruses (N=162) generated along with their complete amino acid sequences.

**Supplemental Table 3.** This table lists all the patient-derived HIV-1 integrase recombinant viruses (N=162) phenotyped, their subtype, the most relevant 27 codons (summarized as “INSTI mutation pattern” in the last column), and fold change (FC) in the 50% effective concentrations (EC50) for bictegravir (BIC), cabotegravir (CAB), dolutegravir (DTG), raltegravir (RAL), and elvitegravir (EVG). n/a denotes no data available.

